# Optic disc centered imaging versus quadrant imaging in retinal oximetry

**DOI:** 10.1101/474122

**Authors:** Ashwin Mohan, Rohit Shetty, Padmamalini Mahendradas, Carroll AB Webers, Tos TJM Berendschot

## Abstract

We aimed to compare intra eye quadrant variation in retinal vessel oxygen saturation in optic disc centered versus quadrant imaging. Forty two consecutive healthy subjects were included in the study. Fifty degree optic disc centered images and images of 4 quadrants (supero-temporal - ST, supero-nasal - SN, infero-nasal - IN and infero-temporal - IT) were taken. The thickest arteriole and venule were chosen in each quadrant in the optic disc centered images. For quadrant images averaged values of 3 segments of thickest arterioles and venules, each above 100µm were chosen. The intra-eye variation between quadrants for arteriolar and venous saturation (%) was compared between optic disc centered and quadrant images. Smaller vessels (70 – 100 µm) in the quadrant images were selected to study the effect of vessel calibre on oxygen saturation. Optic disc centered images gave average arteriolar saturation (%) of 90, 94, 93 and 89 and venous saturation of 58, 60, 60 and 52 in the ST, SN, IN and IT respectively. For quadrant images the average arteriolar saturation was 94, 95, 94 and 91 and the venous saturation was 62, 60, 61 and 60 in the ST, SN, IN and IT respectively. Temporally, the saturation values were significantly different (p<0.001) between optic disc centered and quadrant images. We found no differences nasally. The average intra eye range for arterioles was 11.9 in optic disc centered versus 7.1 in quadrant images (p<0.001) and for venules was 11.6 in optic disc centered versus 7.5 in quadrant images (p<0.001). We found a positive correlation of r=0.18 between saturation and vessel calibre for arteries (p=0.001) and a negative correlation of r=-0.52 for venes (p<0.001). Quadrant imaging significantly reduced intra eye variation between quadrant measurements indicating that maybe the larger variation observed in optic disc centered images are artefactual. Differences in saturation between optic disc centered and quadrant imaging were only seen temporally. We also see physiologically reducing saturation from large arterioles to small arterioles to small venules to large venules.

## Introduction

Retinal oximetry, first described by Hickam *et al*. in 1960 [1], is a technique of measuring relative oxygen saturation in the retinal vessels of the eye. It uses a photospectrometric method with a fundus camera to estimate the saturation. Since then, there have been numerous advancements in retinal oximetry [2] and normative values in various populations have been established. [3–5]

Most of these papers and other published literature use an optic disc centered method of estimation. It has been shown that vessel location within an oximetry image, depending on the gaze of the subject, has a significant effect on measured oxygen saturation in arterioles and venules. [3] Many diseases affect the retina globally and hence global saturations would be sufficient, however many other diseases are local pathologies like chorio-retinitis or vein occlusions and hence it is important to study various angulations of focus.

In our normative database [4] we found that there were large intra-eye differences in the arteriolar saturations with the mean being 94% supero-nasally and 86% infero-temporally. This observation was in agreement with other published normative databases. [5] Paul *et al*. [6] reported that oxygen saturation was lower in veins closer to the optic disc and higher in the superior hemisphere. They also reported that the arteriolar saturations remain relatively constant irrespective of size while the venous saturation are higher in smaller venules.

In our normative database [4] we studied only the largest arteriole or venule per quadrant with an average diameter of 123 µm and 160 µm in arterioles and venules respectively. It would also be interesting to see how the saturation changes with vessel calibre in arterioles and venules.

We thus aimed at comparing the measurements in the same eye in the same setting with optic disc centered versus quadrant images. Since the quadrant images allowed us to study smaller vessels we also studied how the retinal oxygen saturation changed with vessel calibre.

## Methods

### Subjects

Forty two consecutive healthy subjects presenting to our hospital for a routine eye check-up were included in the study. They all had a corrected distance vision acuity more than 20/30, no cataract or other significant media opacities, no history of ocular or systemic disease or history of smoking. The study was approved by the institutional review board and adhered to tenets of the declaration of Helsinki.

### Imaging

The procedure was explained to them in detail and a written informed consent was taken. The subjects were allowed to rest for 5 minutes before the images were taken. None of the subjects had consumed caffeine within 2 hours of the test. After dilatation with 1% tropicamide and 10% phenylephrine all patients had their oximetry images taken on the Oxymap T1 retinal oximeter (Oxymap hf., Reykjavik, Iceland). The right eye was imaged for every subject. Each subject had two 50 degree optic disc centered images and one image in each of 4 quadrants (supero-temporal - ST, supero-nasal - SN, infero-nasal - IN and infero- temporal - IT) taken (see Fig. 1 & 2) – a total of 6 images per eye. The quadrant images were taken such that the edge of the optic disc was at the corner of the image.

**Figure 1.**
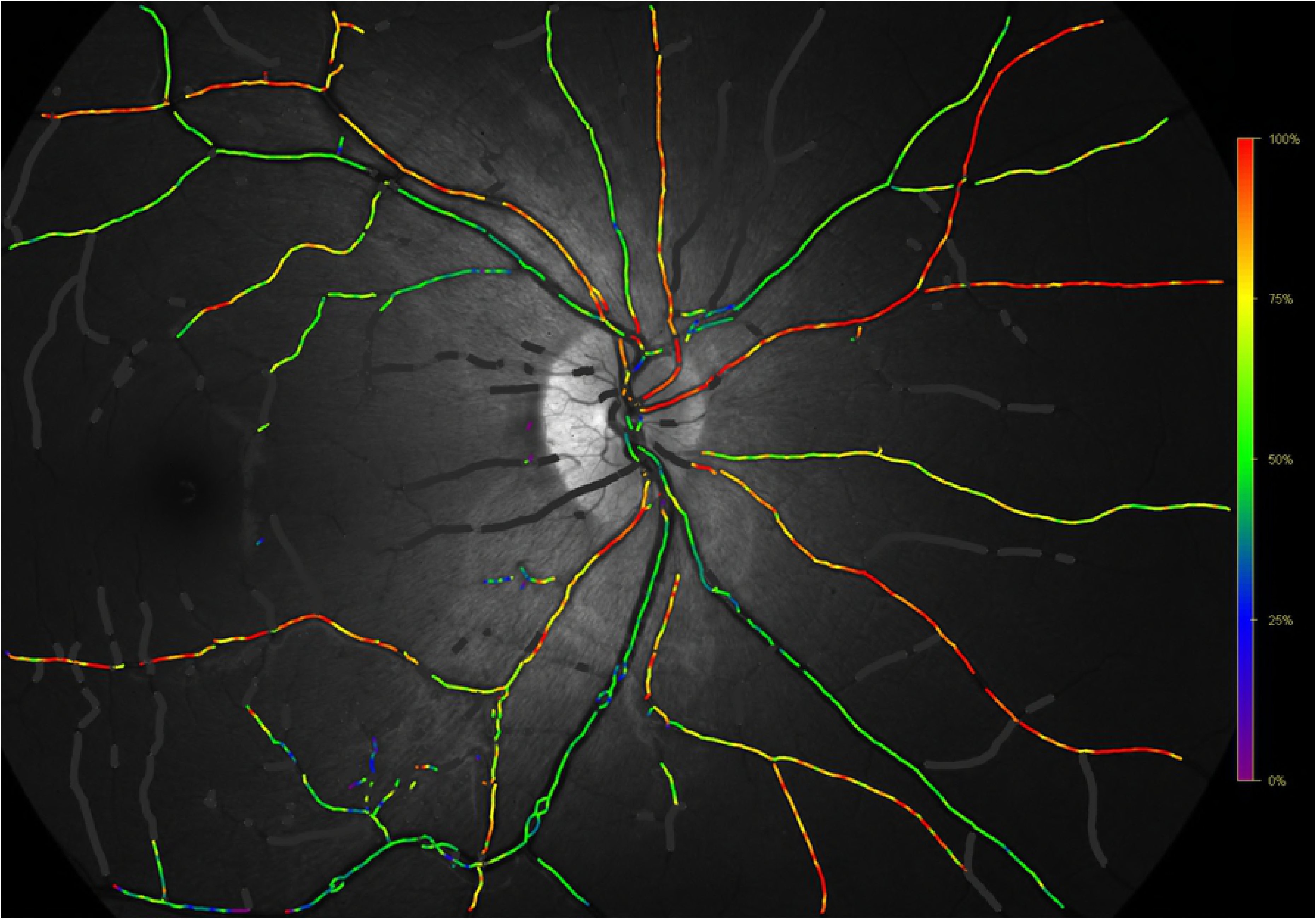
Oximetry maps for the optic disc centered image.

**Figure 2.**
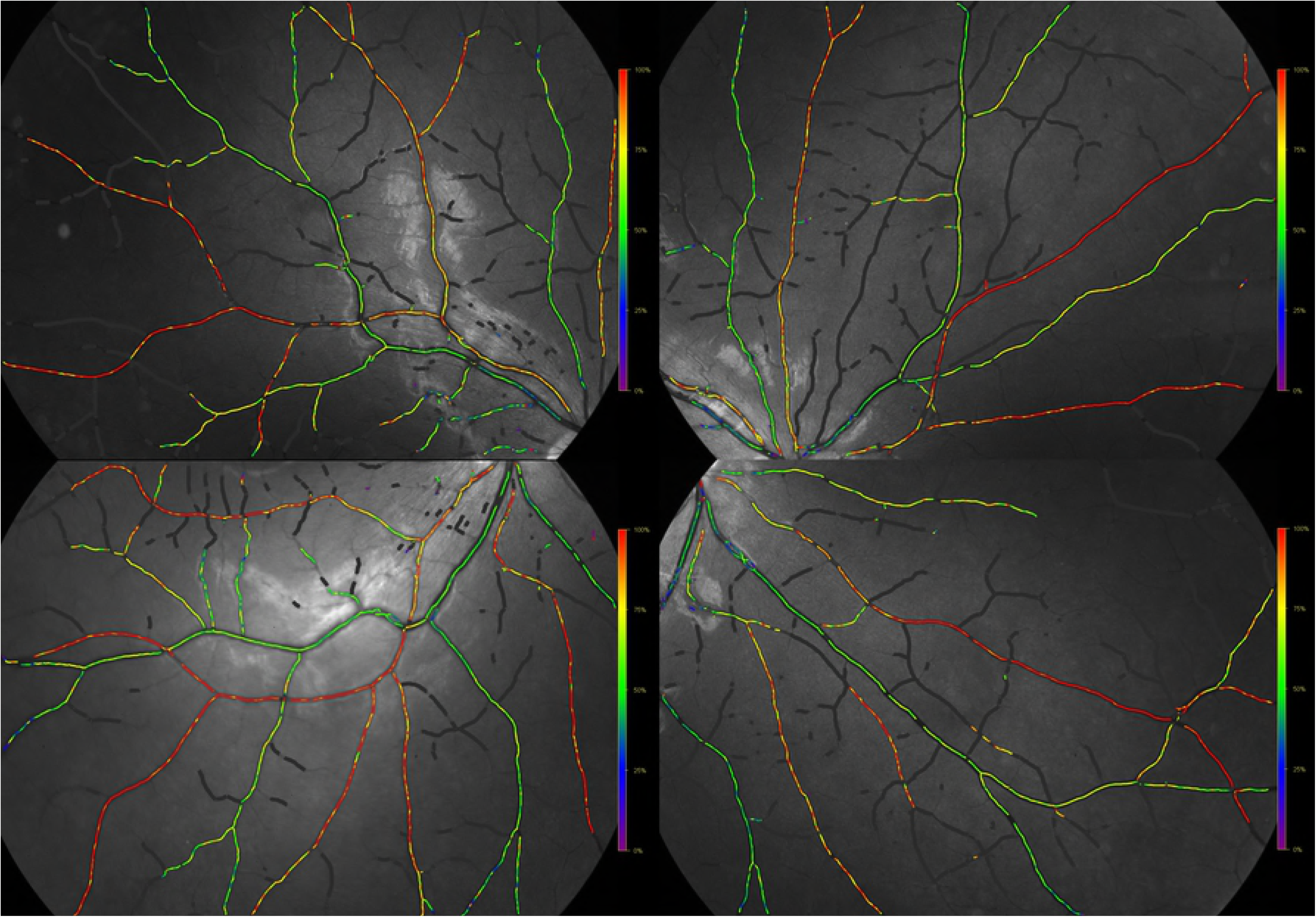
Oximetry maps for the four quadrant images.

### Segment selection

The Oxymap software version used was 2.5.1. Images below 7.5 on the automatic quality analysis were excluded from the analysis. The thickest arteriole and venule were chosen in each quadrant in the optic disc centered images. For quadrant images averaged values of 3 segments of thickest arterioles and venules each above 100 µm were chosen; the segments were between 100 to 150 pixels in length and were selected as much in the centre of the image as possible. The intra-eye range (maximum – minimum) between quadrants for arteriolar and venous saturation (%) was compared between optic disc centered and quadrant images. We also chose smaller vessels (70 – 100 µm) in the quadrant images to study the effect of vessel calibre on oximetry measurements.

### Statistical analysis

All data was entered in Microsoft excel 2013 for Mac and analysed using IBM SPSS v24 for Mac. All values were tested for normality using the Shapiro-Wilk test. The paired t-test was used for comparison of means.

## Results

### Optic disc centered versus quadrant imaging (Table 1 & 2)

**Table 1.** Arteriolar Saturation values in large vessels (>100µm)

|  | ST | SN | IN | IT |
| --- | --- | --- | --- | --- |
| Optic Disc Centered | **$89.6\pm6.4$** | $94.3\pm6.8$ | $93.1\pm8.3$ | **$88.6\pm6.6$** |
| Quadrant | **$93.6\pm3.0$** | $94.7\pm4.1$ | $94.1\pm3.8$ | **$91.1\pm4.3$** |
| p value* | **$<0.001$** | 0.74 | 0.461 | **0.007** |

**Table 2.** Venous Saturation values in large vessels (>100µm)

|  | ST | SN | IN | IT |
| --- | --- | --- | --- | --- |
| Optic Disc Centered | **$57.8\pm6.1$** | $60.4\pm6.4$ | $59.7\pm7.9$ | **$52.1\pm5.9$** |
| Quadrant | **$61.7\pm7.0$** | $60.3\pm6.3$ | $61.2\pm6.2$ | **$59.6\pm5.4$** |
| p value* | **$<0.001$** | 0.893 | 0.242 | **$<0.001$** |

The arteriolar saturation values on optic disc imaging were 89.6±6.4, 94.3±6.8, 93.1±8.3 and 88.6±6.6 and on quadrant imaging were 93.6±3.0, 94.7±4.1, 94.1±3.8 and 91.1±4.3 for supero-temporal (p<0.001), supero-nasal (p=0.740), infero-nasal (p=0.461) and infero- temporal (p=0.007) respectively.

The venular saturation values on optic disc imaging were 57.8±6.1, 60.4±6.4, 59.7±7.9 and 52.1±5.9 and on quadrant imaging were 61.7±7.0, 60.3±6.3, 61.2±6.2 and 59.6±5.4 for supero-temporal (p<0.001), supero-nasal (p=0.893), infero-nasal (p=0.242) and infero- temporal (p<0.001) respectively.

The intra-eye range was 11.9±5.7 in optic disc centered versus 7.1±3.6 in quadrant images for arterioles (p<0.001) and was 11.6±4.5 in optic disc centered versus 7.5±3.2 in quadrant images for venules (p<0.001).

### Within group analysis (Temporal vs Nasal)

The saturations were compared within each vessel group. The temporal saturations were significantly lower than the nasal ones for arterioles in both optic disc (89.1 vs 93.7; p<0.001) and quadrant imaging(92.3 vs 94.4%; p=0.001). For venules they were significantly lower in optic disc centered imaging (55.0 vs 60.1%; p<0.001) however they were comparable in quadrant imaging (60.8 vs 60.7%; p=0.954).

### Effect of vessel calibre

The arteriolar saturations correlated positively with the vessel calibre (r=0.176; p=0.001) (Fig. 3) while the venous saturations correlated negatively with the vessel calibre (r=-0.519; p<0.001) (Fig. 4).

**Figure 3.**
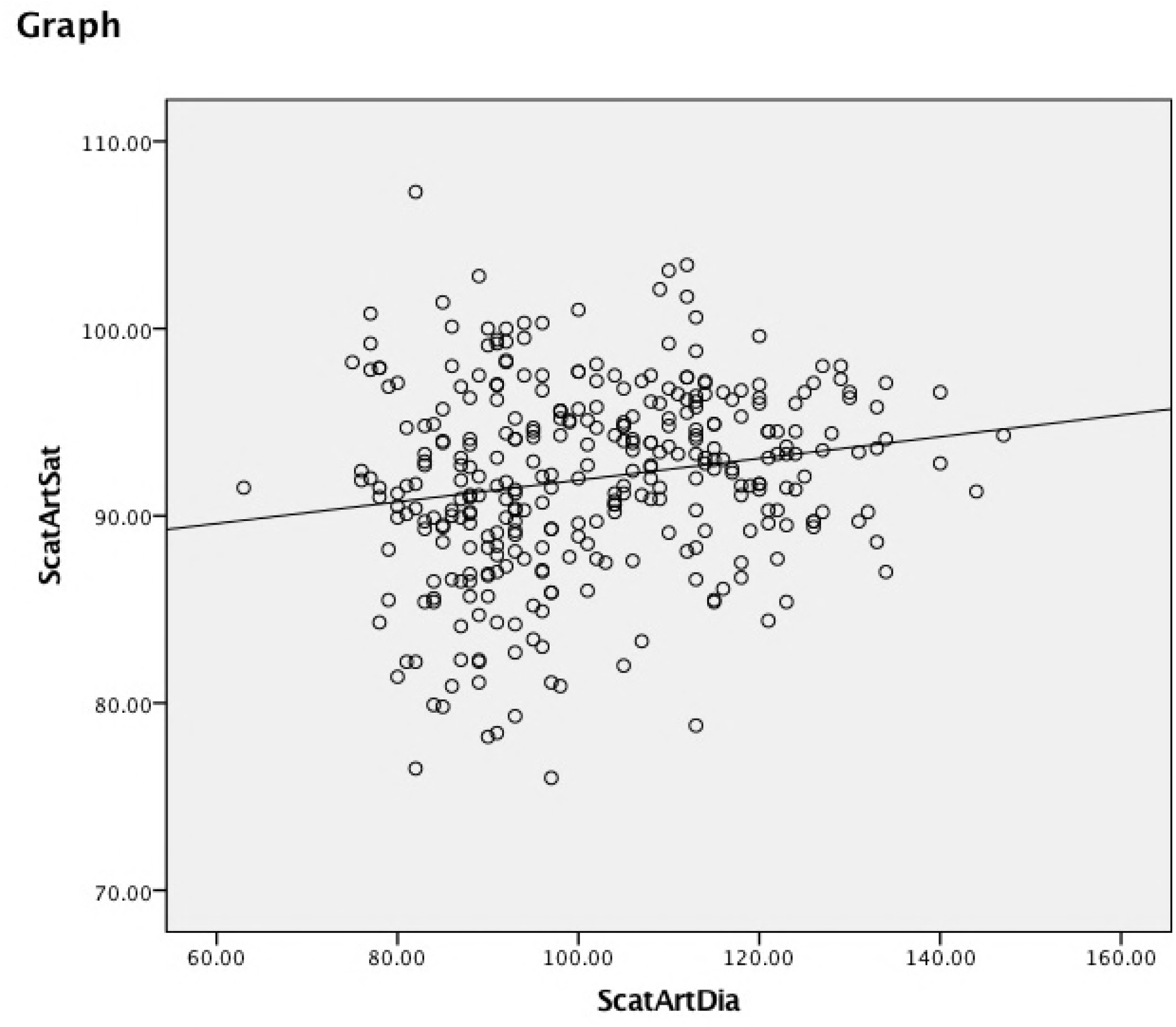
Arterial saturation as a function of arterial diameter. The solid line is a regression line showing the positive correlation (r=0.176; p=0.001).

**Figure 4.**
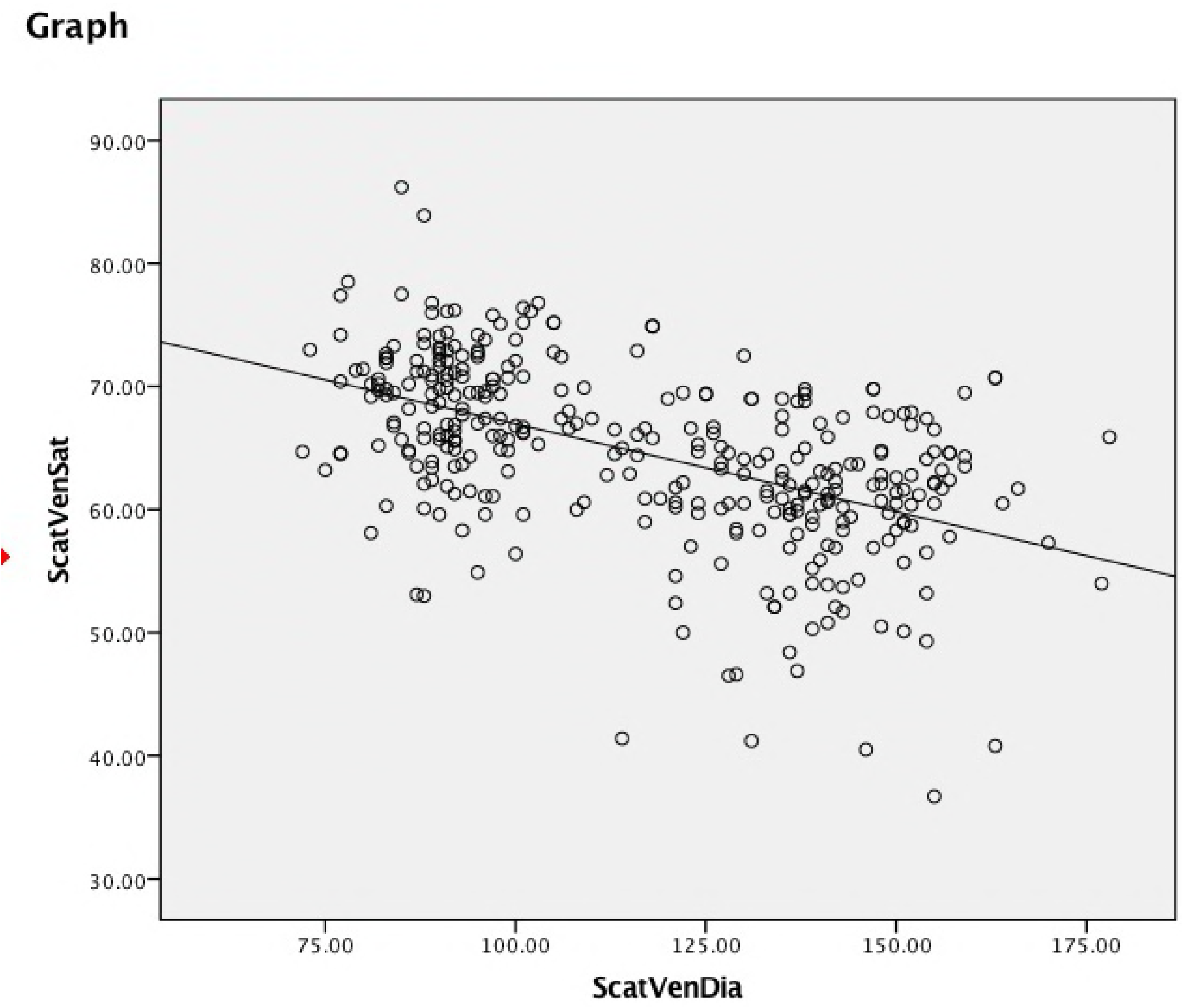
Venous saturation as a function of venous diameter. The solid line is a regression line showing the negative correlation (r=-0.519; p<0.001).

## Discussion

The arteriolar and venous saturation values were significantly different between the optic disc and the quadrant method for measurements made in the temporal hemisphere; they were comparable nasally. The intra eye variation was less in the quadrant imaging as compared to the optic disc imaging. We also saw that in the quadrant imaging the arteriolar saturation decreased with decreasing calibre while it increased in venules.

It is interesting that the difference between the two imaging methods is mainly seen temporally. Our within the group analysis showed that while optic disc centered imaging had significant temporal vs nasal differences for both arterioles and venules the quadrant imaging group had differences only in arterioles. It must also be noted that the difference in arterioles is much lesser in the quadrant imaging versus optic disc centered imaging.

It has been previously established by published normative databases using the optic disc centered method that lower saturations are seen temporally. Jani *et al*. [5] hypothesised that the proximity to the macula could be the cause for the lower saturations. Our study [7] which aimed to study the relationship between perivascular RNFL thickness and oxygen saturations found an inverse relationship. We found that the RNFL was thicker temporally than nasally and hence hypothesised that the thicker RNFL may be extracting more oxygen and hence resulting in the lower saturation. We can argue that if the measured saturations were low due to increased oxygen extraction due to proximity to the macula or due to increased RNFL thickness – i.e. a true decrease in saturations then it must persist even in the quadrant imaging. However we see that this is not true and hence we are led to believe that this could be artefactual.

We also hypothesised in the same study [7] that it may be possible that this could be an optical artefact due to altered background reflectance due to RNFL. The RNFL thickness would remain the same be it optic disc centered or quadrant imaging and hence the effect should persist. This lead us to question if there was some characteristic other than the thickness of RNFL that was influencing reflectance.

An interesting paper by Jeppesen *et al*. [8] report a significant negative association between linear velocity of blood in the vessels and the measured saturations. This can explain why we see lower saturations in the optic disc centered images temporally as the temporal vessels being larger will have a higher linear velocity of blood flow. However this effect disappearing in the quadrant images indicates that linear velocity is only a part of the explanation for the unexplained values seen in retinal oximetry.

Knighton *et al.* [9] performed a study on toad eyes and have established that the reflectance of the RNFL arises from the light scattering of the cylindrical structures of the RNFL. This in turn is dependent on the its orientation relative to the imaging system. In a later study they [10] confirmed that the observed reflectance to a great extent depends on the configuration of the illuminating and viewing apertures and on the retinal position and orientation of the nerve fiber bundles. They found that in optic disc centered images the temporal “ribbons” were brighter than the nasal ones. This apparent brightness can interfere with the background reflectance thus altering measured saturation values in optic disc centered images, since the algorithm used to calculate the saturation assumes a retina based on a limited number of healthy subjects, that ignores and cannot take into account these differences. [2]

We also considered whether the reflectance of the RNFL changes spectrally, thus offering different reflectance at the isosbestic (570nm) and non-isosbestic (600nm) wavelengths. However, two studies by Knighton *et al.* [11,12] showed only small differences between 560 and 680 nm.

Note that in quadrant images each quadrant is centered. Hence the arguments put forward by Knighton *et al.* as described above do not apply and the RNFL is uniformly reflective. This can explain why in this configuration the measurements between the temporal and nasal hemispheres were comparable. It also explains why the intra-eye variation was less in the quadrant images as compared to the optic disc centered images.

The results of the oxy-haemoglobin saturation curve (Fig. 5) offers another argument to support the theory that such large intra-eye variations seen in optic disc centered imaging especially with arterioles must be artefactual. In this curve which is sigmoid in shape we can see that for a haemoglobin saturation (SO_2_) of 97.5% the partial pressure of oxygen (PO_2_) has to be 100 mm Hg, however at a saturation of 80% the PO_2_ drops down to 45%. We can also hypothesise that since the blood comes from the central retinal arteriole and divides into the four major vessels the saturation in the four major vessels should be nearly the same. Each percent difference is saturation requires a change of more than 3 mm Hg in PO_2_; this means that the average 12% difference seen in arteriolar saturations would need a 40 mm Hg difference in PO_2_ which is unexplained physiologically especially in normal eyes. The 12% variation in venules is still more plausible physiologically as firstly the differences in PO_2_ are not as large in the physiological saturation zone of the venules (40-70%) and secondly each major venule would be draining un unequal area of the retina and hence oxygen extraction can differ.

**Figure 5.**
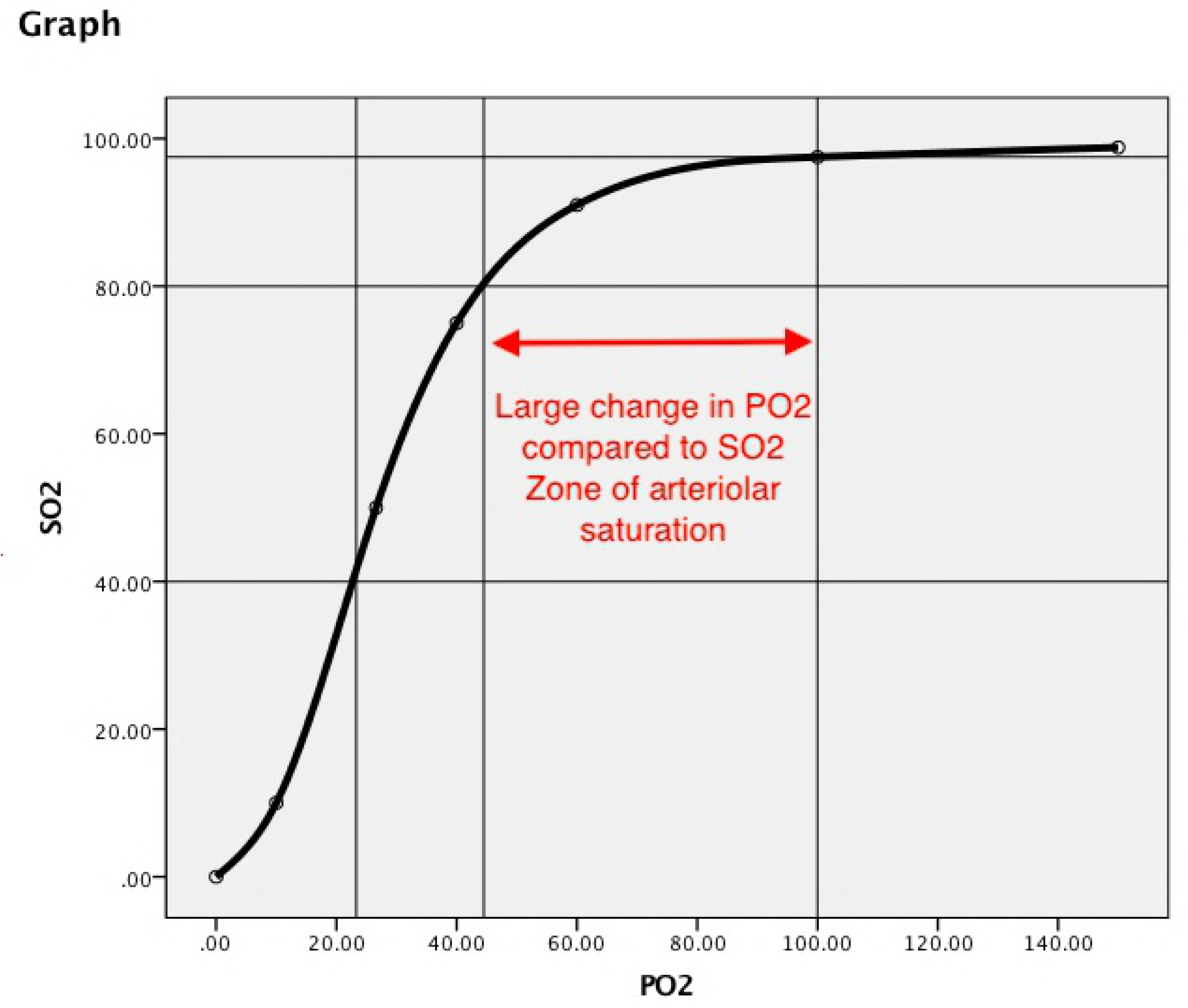
Oxy-haemoglobin saturation curve, showing the haemoglobin saturation (SO_2_) as a function of partial pressure of oxygen (PO_2_)

We noticed that the arteriolar saturation was directly proportional to the calibre whereas the opposite was seen in venules. This trend makes more sense physiologically as blood flows from large arterioles to small arterioles to small venules to large venules. It must be pointed out that this trend is seen in quadrant images whereas our own earlier published normative database[4] with optic disc centered images reported an inverse relationship for both arterioles and venules.

Hence we may conclude that there are significant optical factors that are responsible for the final measured saturation values. These have to be kept in mind when interpreting a retinal oximetry image. Future research should be focused on eliminating these optical artifacts possibly through multispectral imaging or in conjunction with OCT images. The results of this study also indicate that quadrant imaging may be more accurate than optic disc centred imaging which is the standard imaging protocol in most published literature today.

## Aknowledgements

Nil

